## Supplemental information for "Data-driven modeling predicts gene regulatory network dynamics during the differentiation of multipotential progenitors"

### Supplementary Figures

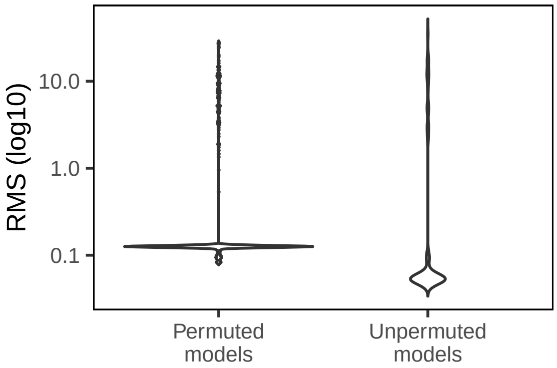


Figure S1: **The significance of gene circuit fits**. The distributions of the RMS scores of gene circuits trained on real data (Unpermuted models) or on randomized synthetic data (Permuted models) are shown as violin plot. The scores were compared using the Wilcoxon ranksum test with continuity correction ($p = 3.8 \times 10^{-8}$).

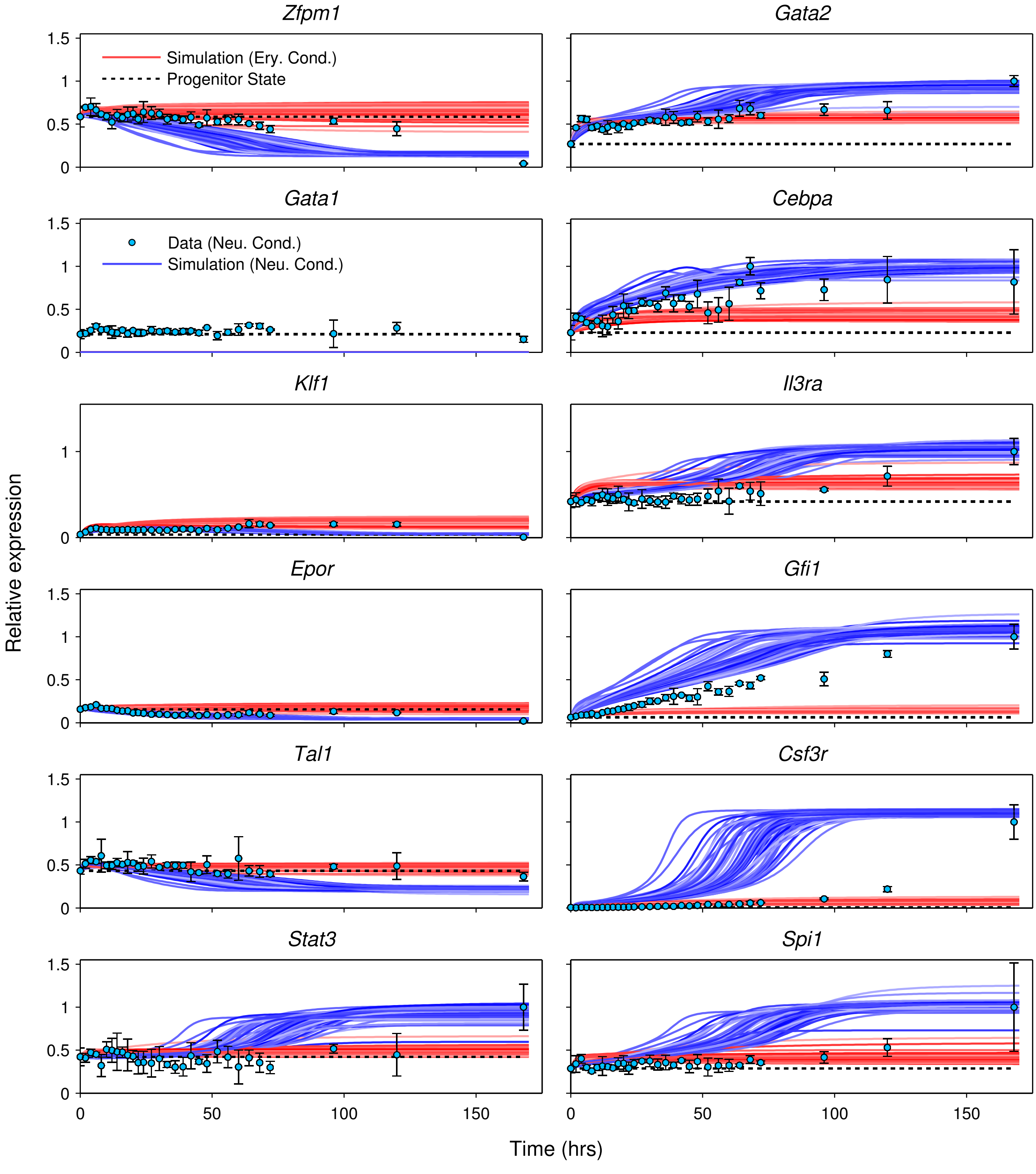


Figure S2: **Simulation of *Gata1* knockout**. *Gata1* knockout was simulated in all 71 models that met the goodness-of-fit criteria. Their output is plotted as lines. The symbols and colors are the same as Figure 1.

### Supplementary Tables

Table S1: **The values of the parameters of the gene circuit models that met the goodness-of-fit criteria.** Columns correspond to parameters while rows correspond to models. $T_{ij}$ are shown as T_Gene$i$_Gene$j$, $b_i$ are shown as b_Gene$i$, $h_i$ are shown as h_Gene$i$, $R_i$ are shown as R_Gene$i$, and $\lambda_i$ are shown as lambda_Gene$i$.

Table S2: **Comparison of model predictions with published experimental evidence.** Each row compares a model prediction about a genetic interconnectivity parameter $T_{ij}$, representing the regulation of gene $i$ by gene $j$, with published experimental evidence. Comparisons of the same parameter to multiple papers are listed in separate rows. $T_{ij}$ is listed as T_Gene$i$_Gene$j$. The prediction column lists the type of regulation inferred by the model. It shows activation or repression when the first quartile of the distribution of the inferred parameter is positive or if the third quartile of the distribution is negative respectively (Fig. 4). The prediction column shows "sign not constrained" when the interquartile range spans negative and positive values. The experiment column lists that type of interaction established in the paper. If the paper describes evidence only of binding but not whether the target is activated or repressed, then the entry is "binding". Negative experimental results are listed as "no effect found". The entry in the experiment column is "not found" if we were not able to find any published tests of the parameter in question. The "status of prediction" columns lists whether the evidence matches the prediction or not. "Confirmed" implies agreements, while "incorrect prediction" implies disagreement. An asterisk indicates that conflicting experimental evidence was found. Conflicting evidence was found for the regulation of *Gata1*, *Spi1*, and *Gata2* by Gfi1 (Moignard *et al.*, 2013, *Nat Cell Biol*, **15**:363–372). Situations where no evidence was found or the paper reported negative results are listed as "undetermined". The "type of evidence" column classifies the evidence as genetic, *cis* regulation, or functional *cis* regulation. Genetic evidence involves genetic manipulation of the predicted regulator followed by a characterization of the target's expression and usually cannot distinguish between direct and indirect effects. *cis* regulatory evidence indicates direct interactions by identifying regulatory elements or binding sites potentially bound by the predicted regulator but does not establish a functional relationship between binding and the expression of the target gene. Functional *cis* regulation goes a step further and manipulates the binding sites and measures reporter or target expression to provide evidence that the binding of the regulator has functional impacts. The organism/cells and citation columns list the organism or cells in which the interaction was tested and the DOI URL for the paper respectively.
